# Multi-omic Analysis Identifies Glioblastoma Dependency on H3K9me3 Methyltransferase Activity

**DOI:** 10.1101/2024.04.18.590173

**Authors:** Qiqi Xie, Yuanning Du, Sugata Ghosh, Saranya Rajendran, Aaron A. Cohen-Gadol, José-Manuel Baizabal, Kenneth P. Nephew, Leng Han, Jia Shen

## Abstract

Histone H3 Lysine 9 dimethylation or trimethylation (H3K9me2 or H3K9me3) marks more than half of the human genome, particularly in heterochromatin regions and specific genes within euchromatic regions. Enzymes catalyzing the methylation of H3K9 have individually been associated with the modulation of gene expression patterns involved in cancer progression, including suppressor of variegation 3-9 homologue 1 (SUV39H1), SUV39H2, SET domain bifurcated 1 (SETDB1), SETDB2, euchromatic histone-lysine N-methyltransferase 1 and 2 (EHMT1/2). However, a comprehensive comparison and understanding of the characteristics and mechanisms of these chromatin-modifying enzymes in cancers remains incompletely understood. In this study, we demonstrated that these six H3K9 methyltransferases differentially expressed in tumors and correlated expression with somatic copy number variations (CNVs) and DNA methylation patterns. Through integrative multi-omics analyses, we identified SUV39H1, SUV39H2, and SETDB1 as the key players among the six H3K9 methyltransferases that exhibited the most significant associations with cancer phenotypes. By incorporating SUV39H1, SUV39H2, and SETDB1, we developed a novel signature termed “H3K9me3 MtSig” (H3K9me3 methyltransferases signature). H3K9me3 MtSig was unique for various tumor types, had prognostic implications and was linked to significant signaling pathways, particularly in glioblastoma (GBM). Furthermore, elevated H3K9me3 MtSig was confirmed in GBM patient-derived cells and tissues. In addition, single-cell expression analysis of H3K9me3 MtSig in GBM tissues demonstrated a pattern related to the G2/M cell cycle and was negatively correlated with immune responses. H3K9me3-mediated repetitive sequence silencing by H3K9me3 MtSig, determined using ChIP-sequencing, contributed to these phenotypes, and inhibiting H3K9me3 MtSig in patient-derived GBM cells suppressed proliferation and increased immune responses. Translationally, H3K9me3 MtSig performed as an independent prognostic factor in a clinical prediction model, and drug susceptibility screening integrating H3K9me3 MtSig identified potential biomarkers and therapeutics for GBM. In summary, H3K9me3 MtSig has the potential to elucidate novel prognostic markers, therapeutic targets, and predictors of treatment response in GBM and other cancer types for clinical intervention.

**Significance:** Differential expressions of H3K9 methyltransferases across cancers correlates with clinical outcomes; in GBM, an H3K9me3 methyltransferase signature links to G2/M cell cycle, immune response pathways, and prognosis, aiding biomarker and treatment development.

## Introduction

Histone H3 lysine 9 dimethylation or trimethylation (H3K9me2 or H3K9me3) is widespread throughout the human genome, notably marking both heterochromatin regions and specific genes within euchromatic regions (1). These regions encompass repetitive DNA sequences such as pericentric satellites, transposable elements, and endogenous retroviruses, along with select euchromatic genes, all commonly subjected to transcriptional silencing. Histone methyltransferases, including suppressor of variegation 3–9 homologue 1 (SUV39H1), SUV39H2, SET domain bifurcated 1 (SETDB1), SETDB2, euchromatic histone-lysine N-methyltransferase 1 and 2 (EHMT1/2), catalyze the methylation of H3K9 and are key players in generating H3K9me3 marks.

SUV39H1, the first identified H3K9 methyltransferase (2), facilitates the establishment and maintenance of heterochromatin structure and transcriptional repression by interacting with heterochromatin protein 1 (HP1). Our previous study demonstrated this process was essential for catalyzing H3K9me3-mediated silencing of repetitive sequences during DNA replication (3). Similarly, SUV39H2 and SETDB1 also catalyze H3K9me3 modifications, albeit with potential differences in expression patterns and regulatory roles in specific cellular contexts (4,5). Although SETDB2 shares the ability to methylate H3K9me3, its specific functional implications remain largely unexplored compared to SETDB1 (6,7). EHMT1 and EHMT2, highly homologous proteins, form a heteromeric complex that primarily facilitates H3K9 monomethylation (H3K9me1) and H3K9me2 modifications to regulate gene expression (8).

Dysregulation of H3K9 methyltransferases leads to abnormal H3K9 methylation patterns, impacting gene expression and chromatin dynamics and contributing to various diseases, including cancer. Previous research indicates that SETDB1 plays a role in suppressing endogenous retroelements to evade immune surveillance and promote tumor development in melanoma (9). Another study suggests that the methyltransferase complex containing SETDB1 and SUV39H1 represses developmental genes, such as *Hox* genes, resulting in alterations of gene expression that accelerate melanoma (10). In breast cancer, SUV39H1 deposits H3K9me3 on the *E-cadherin* gene, suppressing its expression, potentially driving epithelial to mesenchymal transition (EMT) and metastasis (11). Additionally, EHMT2 suppresses the expression of the cell adhesion molecule Ep-CAM, promoting EMT and tumor progression in lung cancer (12).

H3K9 methyltransferases have individually been strongly implicated in tumorigenesis and progression. However, the genomic and clinical attributes of H3K9 methyltransferases in tumors and their synergistic functions and regulatory molecular mechanisms remains incomplete. To investigate the functional significance of H3K9 methyltransferases in tumors, we carried out an integrated multi-omic analyses, combining genomic, transcriptomic, epigenomic, and clinical data, together with rigorous experimental validation.

## Materials and Methods

### Datasets, gene expression analysis, and survival analysis

In the mRNA differential expression analysis, normalization was applied to the mRNA expression of 20 cancer types with paired data from The Cancer Genome Atlas (TCGA). Fold change was represented by mean (Tumor)/mean (Normal), and P-values were determined using t-tests, with adjustment through False Discovery Rate (FDR). Genes with log2(fold change) > 1 and FDR < 0.05 were retained for further analysis. Survival analysis combined mRNA expression data of the H3K9 methyltransferases and corresponding clinical survival data across 33 cancer types for expression survival analysis. Functional similarity was evaluated using the geometric mean of semantic similarity in Molecular Function (MF) and Cellular Component (CC) (13). Proteins within interaction groups are functionally and spatially related, forming a crucial aspect of their functional role. Proteins within the H3K9 methyltransferases interaction group were ranked based on the average functional similarity, with a cutoff value of 0.60. A further analysis of the correlation of the H3K9 methyltransferases across all cancers was conducted. To explore the relationship between the expression of the H3K9 methyltransferases and the objective response rate (ORR) to anti-PD-1/PD-L1 therapy (14,15), tumor samples were stratified into high and low groups based on gene medians. Cox regression models were employed to fit survival time and status for both groups, calculating the hazard ratio (HR) for each gene. The results from the Cox regression model were used to assess the correlation of H3K9 methyltransferases gene expression with pan-cancer overall survival (OS), disease-specific survival (DSS), disease-free interval (DFI), and progression-free interval (PFI).

### Single nucleotide variation analysis, copy number variation analysis and methylation analysis

Genomic positions of the H3K9 methyltransferases were depicted in a circular plot. Single-nucleotide polymorphism (SNV) data, copy number variation (CNV) data, and methylation data for 33 cancer types were collected from TCGA database (7). Initially, all 33 cancer types were included in the pan-cancer analysis (Table S1). After excluding cancers lacking paired normal tissue data, the subsequent analysis included 20 cancer types, namely BLCA, BRCA, CESC, CHOL, COAD, ESCA, GBM, HNSC, KICH, KIRC, KIRP, LIHC, LUAD, LUSC, PAAD, PRAD, READ, STAD, THCA, and UCEC.

For the 33 cancer types, SNV data were calculated using the formula: mutation samples/total cancer samples, and the R software maftools package (16) was used to generate the SNV cancer gene plot (oncoplot). The CNV and methylation analyses incorporated paired data from 20 cancer types, enabling comprehensive examination within this cohort. CNV was categorized into two subtypes: homologous and heterologous, encompassing amplifications and deletions, to represent the occurrence of CNV on one or both chromosomes. Percentage statistics were generated based on CNV subtypes, merging the mRNA expression of the H3K9 methyltransferases with raw CNV data. The P-values were adjusted using FDR to explore the correlation between them.

For methylation data, t-test was conducted for methylation differences between tumor and normal samples and P-values were corrected with FDR. FDR < 0.05 was considered significant. The mRNA expression and methylation data of the H3K9 methyltransferases were merged, the correlation between paired mRNA expression and methylation was examined using the Spearman correlation coefficient. P-values were FDR-corrected, and genes with FDR < 0.05 were retained. All plots were created using the R software ComplexHeatmap package (17).

### Evaluation of pan-cancer prognosis linked to H3K9me3 methyltransferases signature (H3K9me3 MtSig)

The R software GSVA package (18) for single-sample gene set enrichment analysis (ssGSEA) was used to calculate the enrichment scores (ES) of SUV39H1, SUV39H2, and SETDB1, defined as the H3K9me3 MtSig score. Based on the H3K9me3 MtSig score-associated genes, a Cox proportional hazards regression model was established. If the coefficient β was obtained for each gene associated with H3K9me3 MtSig, each patient would receive an H3K9me3 MtSig as follows. Univariate Cox regression analysis evaluated the correlation between H3K9me3 MtSig expression and OS, DSS, and PFI. The optimal value was used as the threshold to categorize patients into high and low expression groups. Survival differences between high and low groups were analyzed for various cancers. P-values were calculated through the log-rank test, and P < 0.05 was considered statistically significant.

### Analysis of signaling pathways associated with H3K9me3 MtSig

Pan-cancer and tumor type samples were stratified into top and bottom 30% based on H3K9me3 MtSig expression. Correlation analysis using GSEA explored associations between H3K9me3 MtSig and all hallmark pathways, with the top 3 gene sets applied in the z-score algorithm implemented in the R software GSVA package. Correlation analyses were performed between H3K9me3 MtSig and the G2/M checkpoint, as well as immune responses pathways, in each cancer type. Furthermore, correlations between SUV39H1, SUV39H2, SETDB1, with these pathways were examined. For GBM analysis, the Extended GBmap dataset (bioRxiv 2022.08.27.505439), comprising 16 datasets gathering transcriptional profiles from 110 GBM patients and spanning over 330,000 cells, was utilized to analyze the single-cell transcriptional data of H3K9me3 MtSig, G2/M checkpoint, and immune responses. The Extended GBmap dataset integrates and re-analyzes multiple published single-cell RNA-seq datasets for GBM, providing a comprehensive resource for exploring cellular heterogeneity and gene expression patterns across diverse GBM samples. Specifically, the pre-processed and integrated gene expression matrices provided in the Extended GBmap resource were extracted and analyzed for expression profiles of genes associated with the H3K9me3 MtSig, G2/M checkpoint, and immune responses pathways. The integrated dataset enabled us to perform comparative analyses and identify associations between the expression patterns of these gene signatures and cellular states within the GBM tumor microenvironment. Additionally, the correlations between SUV39H1, SUV39H2, SETDB1, and key molecules of the G2/M checkpoint pathway, namely CDK1 and MKI67, and immune responses pathway, namely CCL7, CXCL8, CD274 and PDCD1 were analyzed based on the data from TCGA and Chinese Glioma Genome Atlas (CGGA) databases.

### ChIP and ChIP-seq analysis

The H3K9me3 ChIP experiment followed our previously reported protocol (3). Briefly, 10^6^ GSC3565 cells, which originated from a GBM stem cell (GSC) model established using cells obtained from a 32-year-old male GBM patient at University Hospitals, Cleveland, in Dr. Jeremy Rich’s lab (19), underwent cold PBS washes, and native chromatin was extracted using a commercial kit (Abcam, cat# ab117152). Chromatin was re-suspended in ChIP lysis buffer and sonicatedin a S220 Focused-Ultrasonicator (Covaris) for 7 min (Duty cycle-5%, Intensity-4, Cycles/Burst-200). Following centrifugation, the supernatant was diluted and subjected to H3K9me3 ChIP using the ChIPAb+ Trimethyl-Histone H3 (Lys9) kit (Millipore, cat# 17-10242). Bound material was recovered after incubation with 15 μL of protein G beads (Millipore, cat # 16-266) at 4°C, washed sequentially, and eluted. DNA was extracted using the Gel Extraction Kit (Qiagen cat# 28704) and submitted to Novogene for standard sequencing. For data analysis, ChIP-seq reads were aligned to the Homo sapiens genome (hg38) using Bowtie 2 (20) with default parameters, which allows multiple alignment. Peaks were called using MACS2 software (21) with default parameters. The midpoint of each estimated fragment by MACS2 and its location on the genome was calculated. The genome was divided into non-overlapping windows of the default 100 bp. An aligned read was considered to be located in a window if the midpoint of its estimated fragment was within the window. The number of midpoints in each window was counted and an empirical distribution of window counts was created. A zero-truncated negative binomial model was fit to the distribution, and a peak was determined based on the FDR (0.001, default) calculated from the model. Overlapping enriched windows were merged into regions and reported. For analysis of REs, a list of H3K9me3 peaks associated with REs was created by intersecting H3K9me3 binding peaks with RE loci obtained from RepeatMasker (https://github.com/rmhubley/RepeatMasker), which was used in the downstream analysis. Reads were mapped and assigned to the H3K9me3-associated REs using EDTA with the recommended parameters (https://github.com/oushujun/EDTA). The resulting counts for REs were analyzed by the edgeR package (22) to obtain CPM values. The ChIP-seq data of D456 and GSC23 cells were obtained from GEO: GSE119081 (23). The BigWig files were obtained using deeptools (24). The heatmap and profile plots for ChIP-seq data were performed by deeptools plotHeatmap and plotProfile in the scale-regions mode. Venn diagrams were obtained using VennDiagram package (25). The ChIP-seq binding signal from BAM files was visualized in The Integrative Genomics Viewer (IGV) (26).

### Clinical prediction model

Conditional survival analysis was conducted on GBM survival data from TCGA, followed by evaluation of the associated risk scores for each GBM patient. Based on the H3K9me3 MtSig, GBM patients were stratified into low H3K9me3 MtSig score group (< median) or high H3K9me3 MtSig score group (≥ median), and a risk factor association analysis was performed. To develop a prognostic line chart predicting GBM survival rate (27), age and stage were used to predict the overall survival of GBM patients. All P-values were based on two-sided tests, and P < 0.05 was considered statistically significant. To ensure the credibility of clinical prediction model, decision curves and calibration curves were employed for model validation (28). To further ensure the generalizability of the model, additional validation using GBM data from CGGA was performed.

### Drug susceptibility screening

The expression profile data for human cancer cell lines were obtained from the Broad Institute Cancer Cell Line Encyclopedia (CCLE) project (https://portals.broadinstitute.org/ccle/) (29). The drug sensitivity data for human cancer cell lines were derived from the Cancer Therapeutics Response Portal (CTRP; https://portals.broadinstitute.org/ctrp) (30) and the PRISM repurposed dataset (https://depmap.org/portal/prism/) (31). The CTRP contains sensitivity data for 481 compounds across more than 860 human cancer cell lines, while the PRISM dataset includes sensitivity data for 1,448 compounds across more than 482 human cancer cell lines. Both datasets provide areas under the dose-response curve (AUC) values as a measure of drug sensitivity, with lower AUC values indicating increased sensitivity to treatment. Missing AUC values were imputed using the K-nearest neighbors (K-NN) interpolation method. The databases allow for the selection of drugs based on genomic features of cancer patients. The drug set evaluation focused on assessing the sensitivity of compounds to GBMs with high H3K9me3 MtSig.

### Western blot

Normal brain stem cells (NSC 194 (19)) and patient-derived GBM cells (GSC1914 (19), GSC3565 (19), GSC839 (32), and GSC2907 (32)) were generously provided by Dr. Jeremy Rich from the UPMC Hillman Cancer Center, Pittsburgh, PA, USA. These cells were cultured using neurobasal media (Gibco, cat# 12349015) supplemented with B27 lacking vitamin A (Gibco, cat# 12587010), 20 ng/ml EGF (R&D Systems, cat# 236-EG-01M), 20 ng/ml bFGF (R&D Systems, cat# 3718-FB-025), 1% sodium pyruvate (Gibco, cat# 11360070), 1% GlutaMAX (Gibco, cat# 35050061), and 1% penicillin/streptomycin (Cytiva, cat# SV30010). The cells were lysed in cold RIPA buffer (Thermo Fisher Scientific, cat# 89901) supplemented with Protease and Phosphatase Inhibitor Mini Tablets (Thermo Fisher Scientific, cat# A32961). After brief sonication and clarification, lysates underwent SDS-PAGE and transferred to Immun-Blot PVDF Membrane (Bio-Rad, cat# 1620177). Following blocking with 5% non-fat milk TBS containing 0.1% Tween-20 (TBST), membranes were incubated overnight at 4°C with primary antibody. Subsequently, membranes were washed and probed with species-specific HRP-conjugated secondary antibodies. After washing, membranes were developed using Clarity Max Western ECL Substrate (Bio-Rad, cat# 1705062) before imaging with a ChemiDoc Imaging System (Bio-Rad). Antibodies used included anti-SUV39H1 (Thermo Fisher Scientific, cat# 702443, 1:1000), anti-SUV39H2 (Proteintech, cat# 11338-1-AP, 1:300), anti-SETDB1 (Proteintech, Cat# 11231-1-AP, 1:1000), and anti-α-Tubulin (Proteintech, Cat# 11224-1-AP, 1:4000).

### Immunohistochemistry

Normal brain and GBM tissues were obtained from the Enterprise Clinical Research Operations (ECRO) Biorepository at Indiana University Health Methodist Hospital with approval from the Institutional Review Board (IRB) for the collection of human biological materials. Detailed information regarding the tissues can be found in Table S2. Subsequently, these tissues were processed and preserved in formalin-fixed paraffin-embedded (FFPE) format, deparaffinized in xylene and rehydrated through a graded ethanol dilution. Antigen retrieval was done through heat induction using appropriate retrieval buffer. Endogenous peroxidase activity was reduced using endogenous peroxidase and alkaline phosphatase blocking solution (Blockxol, cat# SP-6000) followed by blocking for 1 hr at room temperature using animal-free blocker (Vector Laboratories, cat# SP-5035). Then the sections were stained with primary antibody diluted in SignalStain Antibody Diluent (Cell Signaling Technology, cat# 8112L) overnight at 4°C. After washing, the sections were incubated with SignalStain Boost Detection Reagent (Cell Signaling Technology, cat# 8114S) of correct species for 30 mins at room temperature. DAB (3,3′-Diaminobenzidine, Cell Signaling Technology, cat#11725P) was applied to each section, and once the sections developed, they were rinsed in water and counterstained with hematoxylin. After dehydration, slides were mounted using organic mounting media (Vector Laboratories, cat# H-1700). The slides were imaged using Motic EasyScan Pro 6 (Motic). The antibody used was anti-SUV39H1 (Thermo Fisher Scientific, cat# PA5-29470, 1:100), anti-SUV39H2 (Thermo Fisher Scientific, cat# MA5-34753, 1:50), and anti-SETDB1 (Thermo Fisher Scientific, cat# 11231-1-AP, 1:200).

### Immunofluorescence

GSC3565 cells cultured on Matrigel-coated coverslips (Corning, cat# 356231) were transfected with GFP-H3.3 or GFP-H3K9M plasmids using FuGENE HD Transfection Reagent (Promega, cat# E2311). After 48 hrs, the cells were washed with PBS, fixed with 4% paraformaldehyde for 20 mins, and permeabilized with 0.1% Triton X-100 in PBS for 10 mins at room temperature. After blocking with 1% BSA in PBS for 1 hr, cells were incubated with anti-H3K9me3 primary antibody (Cell Signaling Technology, cat# 13969, 1:500) overnight at 4°C. Subsequently, cells were stained with Alexa Fluor 594-labeled anti-Rabbit secondary antibody (Invitrogen, cat# A11012, 1:500) for 1 hr, mounted with anti-fade mounting medium containing DAPI (Vector Laboratories, cat# H-1200) for 5 mins, and visualized using fluorescence microscopy (Nikon DS-Fi3).

### qPCR analyses

Total RNA from the GSC3565 cells was isolated using the RNeasy Plus Mini Kit (QIAGEN, cat# 74136). cDNA was synthesized using qScript cDNA SuperMix (Quantabio, cat# 95048) following the manufacturer’s instructions. qPCR reactions were performed using PowerUp SYBR Green Master Mix (Applied Biosystems, cat# 25742). Thermal cycling conditions included an initial denaturation step of 95 °C for 30s, 40 cycles at 95 °C for 3 s, and 60 °C for 20 s. The qPCR primers used for gene expression analysis included: mcBox-AGGGAATGTCTTCCCATAAAAACT and GTCTACCTTTTATTTGAATTCCCG, IFNα- AATGACAGAATTCATGAAAGCGT and GGAGGTTGTCAGAGCAGA, CDK16- TCCGTCGTGTCAGCCTATCT and TCATGTTCCAGTCTGATCTCCTT, and Actin-CATGTACGTTGCTATCCAGGC and CTCCTTAATGTCACGCACGAT.

## Results

### Expression patterns and prognostic relevance of H3K9 methyltransferases

We interrogated the expression levels of the H3K9 methyltransferases in a variety of tumors. It was observed that, across various cancers, except for EHMT1 and SETDB2, which exhibited relatively low expression levels, the expression levels of other H3K9 methyltransferases were generally upregulated with log2 fold changes greater than 1, while they were downregulated by 40% to 60% in KICH, KIRP, PAAD, and THCA (Fig. 1A). SUV39H1 showed a high similarity in expression patterns with SUV39H2, SETDB1, and EHMT2 across tumors, whereas SETDB2 exhibited relatively similar and predominantly low expression patterns with EHMT1 (Fig. 1A). Functional similarity analysis of these six H3K9 methyltransferases, based on the geometric mean of semantic similarities in Molecular Function (MF) and Cellular Component (CC) annotations, revealed that SUV39H2 and SUV39H1 might play central roles with SUV39H2 being the highest critical value score among them (Fig. 1B). Using Spearman correlation analysis, we observed a strong correlation in the expression of H3K9 methyltransferases across various cancers, particularly among SUV39H1, SUV39H2, and SETDB1 (Fig. 1C). Survival analysis showed that in GBM, SARC, LIHC, and BRCA, the high expression levels (top 25%) of the H3K9 methyltransferases were primarily associated with poorer survival with hazard ratios of 1.5 to 2.2 (Fig. 1D). Conversely, in THYM and READ, the H3K9 methyltransferases mainly functioned as protective factors linked to better survival (P < 0.05, Fig. 1D). Furthermore, when analyzing the correlation of H3K9 methyltransferases with OS, DSS, DFI, and PFI, we found SUV39H1, SUV39H2, and SETDB1 revealed similar results and their high expression levels were associated with poorer prognosis with hazard ratios of 1.3 to 1.8 in GBM and KIRP tumors (Fig. S1A - F). The analysis of H3K9 methyltransferases with the ORR to anti-PD-1/PD-L1 checkpoint blockade immunotherapy revealed correlations with SUV39H1, SUV39H2, and SETDB1 (Fig. 1E - G). Notably, a positive correlation was observed between SUV39H1 expression and ORR (P < 0.05), suggesting that elevated levels of SUV39H1 are linked to improved response rates across various tumors. These findings demonstrate heterogeneous expression patterns of H3K9 methyltransferases across cancers, with some members correlating with expression, and that the expression of these enzymes correlates with cancer prognosis and patient outcomes.

**Fig. 1.**
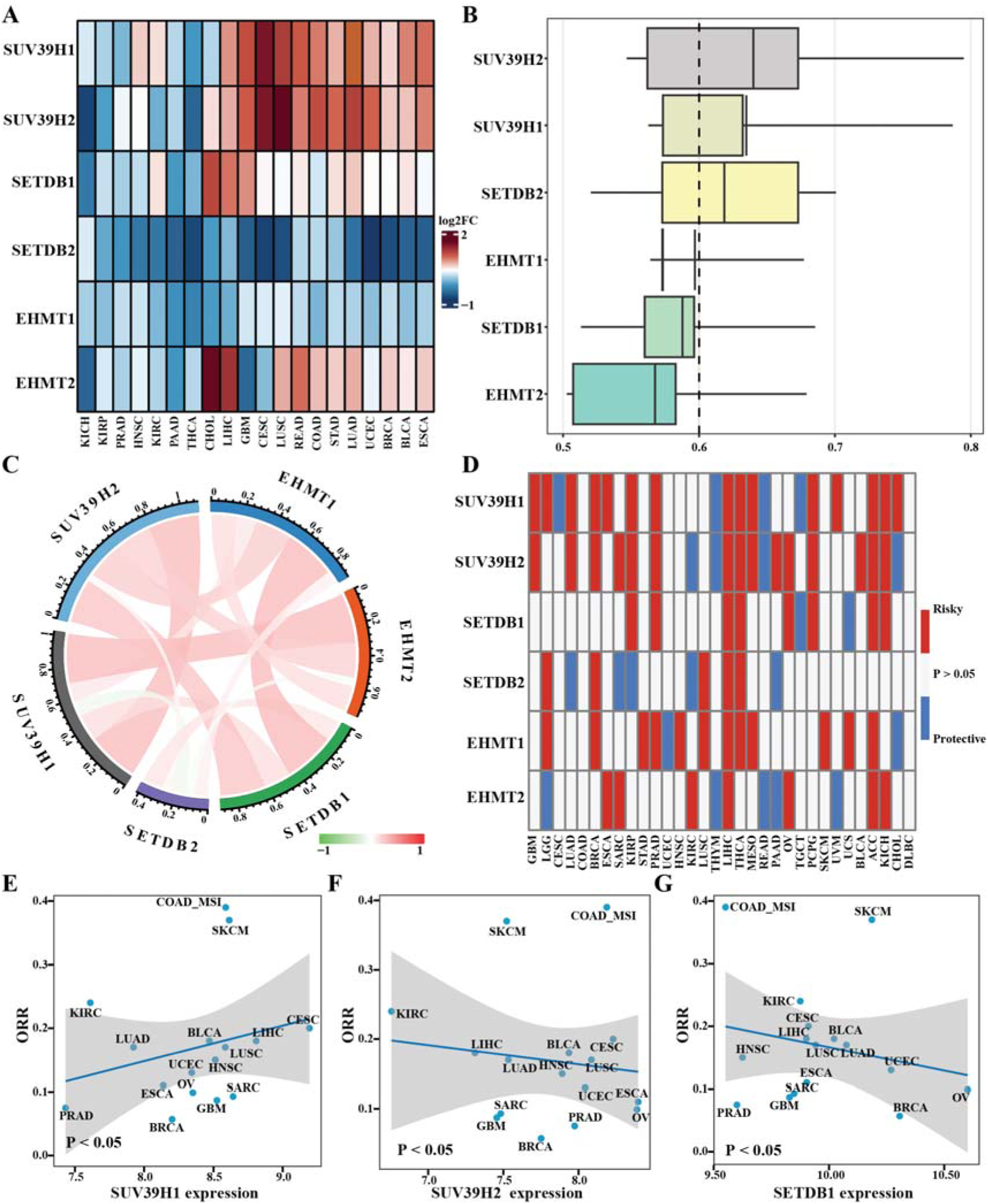
Expression patterns and prognostic relevance of H3K9 methyltransferases in pan-cancer. (A) The expressions of the six H3K9 methyltransferases in pan-cancer, with red indicating upregulation and blue indicating downregulation. (B) Functional similarity analysis of H3K9 methyltransferases in a boxplot. The box represents the middle 50% of similarity, while the upper and lower bounds indicate the 75th and 25th percentiles of similarity. The line within the box represents the mean functional similarity. Proteins with high average functional similarity (cut-off > 0.60) are considered central proteins in the granulation enzyme interaction network. The dashed line represents the cut-off value. (C) Expression correlations of H3K9 methyltransferases in pan-cancer. (D) The prognostic value of the H3K9 methyltransferases was analyzed using univariate Cox regression, where red represents a correlation between high expression of H3K9 methyltransferases and poor survival, and blue represents a correlation with better survival. (E to G) Analysis of SUV39H1 (E), SUV39H2 (F), SETDB1 (G) with the objective response rate (ORR) to anti-PD-1/PD-L1 immunotherapies.

### Genetic and epigenetic alterations of H3K9 methyltransferases genes

Analysis of the genomic locations of H3K9 methyltransferase genes revealed that *SUV39H1* is situated on the X chromosome, *SUV39H2* and *EHMT1* on chromosome 9, *SETDB1* on chromosome 1, *SETDB2* on chromosome 13, and *EHMT2* on chromosome 6 (Fig. 2A). To discern dysregulation patterns in H3K9 methyltransferases, we analyzed genomic and epigenomic data encompassing genetic variations, CNVs, mRNA expression, and DNA methylation across 20 cancer types and normal tissues. We observed high mutation frequencies ranging from 19% to 25% for SETDB1, EHMT1, and EHMT2 across multiple cancer types (Fig. 2B). Examination of CNV percentages for H3K9 methyltransferase genes revealed high frequencies (greater than 5% in all samples) in most cancer types, except for THCA, where CNV frequencies were lower at around 1% to 3%, possibly due to the overall low CNV frequency in THCA. The H3K9 methyltransferase genes displayed distinct CNV patterns, with similar frequencies of 6% to 12% observed for *SUV39H1* and *SUV39H2* across major cancer types (Fig. 2C). We conducted an analysis of mRNA expression and CNV data for H3K9 methyltransferase genes from TCGA. The findings indicated a generally positive correlation with Spearman’s r values of 0.6 to 0.8 between mRNA expression and CNV, particularly notable for *SETDB1*, *SETDB2,* and *EHMT1* in BLCA, BRCA, LUAD, and LUSC (P < 0.05, Fig. 2D). Conversely, SUV39H1 displayed an opposing pattern to other H3K9 methyltransferase genes in most tumors.

**Fig. 2.**
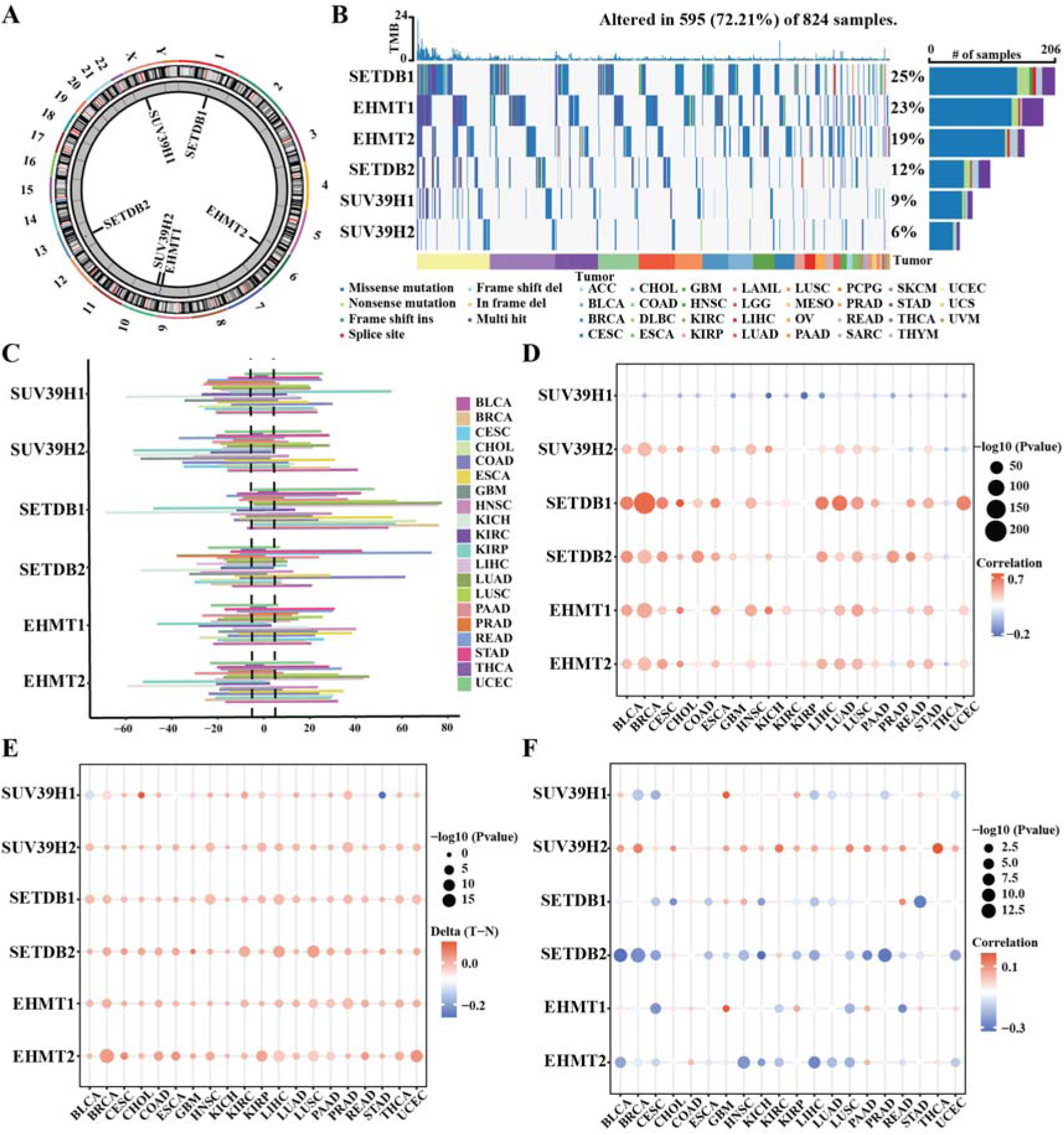
Genetic and epigenetic alterations of the H3K9 methyltransferases genes. (A) Circular plot showing the gene positions of the H3K9 methyltransferases. (B) Waterfall plot demonstrating the mutation frequency of the H3K9 methyltransferase genes in pan-cancer. (C) Histogram showing the frequency of CNVs for H3K9 methyltransferase genes in pan-cancer. (D) Spearman correlation analysis of the transcriptional expression and CNVs of the H3K9 methyltransferase genes. (E) DNA methylation in the H3K9 methyltransferase genes in pan-cancer. Genes with high and low DNA methylation are marked in red and blue, respectively. (F) Spearman correlation analysis of DNA methylation and expression of the H3K9 methyltransferase genes.

To elucidate epigenetic regulation of H3K9 methyltransferase genes, we examined the DNA methylation status. The data revealed a high degree of DNA methylation consistency across different tumors, with most cancers exhibiting significant DNA methylation of H3K9 methyltransferase genes (P < 0.05). Interestingly, *SUV39H1* exhibits a pattern opposite to other H3K9 methyltransferase genes in BLCA and STAD (P < 0.05, Fig. 2E). Correlation analysis between DNA methylation and mRNA expression demonstrated a general negative correlation for most genes, except for *SUV39H2*, which exhibited a positive correlation (P < 0.05, Fig. 2F). These findings suggest that mutations, CNV, and DNA methylation regulate expression of H3K9 methyltransferase genes, impacting tumorigenesis and progression.

### H3K9me3 methyltransferases signature (H3K9me3 MtSig) and regulated pathways in cancers

The expression and survival analysis findings suggest that SUV39H1, SUV39H2, and SETDB1, all primarily involved in methylation of H3K9me3, exhibit similar patterns and correlations in cancers. We integrated these three H3K9 methyltransferases into a unified signature named H3K9me3 methyltransferases signature (H3K9me3 MtSig). We compared the differential expression of H3K9me3 MtSig between tumor and normal tissues in TCGA dataset (Fig. 3A). In the majority of tumors, H3K9me3 MtSig was significantly upregulated compared to the normal, except for THCA (Fig. 3A). Patients in the TCGA training cohort were stratified into high-risk and low-risk groups based on whether their H3K9me3 MtSig exceeded the population median. High H3K9me3 MtSig was linked to adverse survival outcomes, including decreased DSS, OS, and PFI, compared to the low-risk group (Fig. 3B). Using GSEA on transcriptome data from tumors exhibiting the highest and lowest 30% H3K9me3 MtSig levels, we explored the cell signaling pathways associated with H3K9me3 MtSig. H3K9me3 MtSig displayed a notable positive correlation with G2/M checkpoint pathway (Fig. 3C), indicative of cancer cell proliferation. Conversely, it exhibited a negative correlation with the immune response pathways, including inflammatory response, interferon alpha response, and interferon gamma response (Fig. 3D). Additionally, hallmark pathway analysis revealed a positive correlation (R=0.63, P < 0.001) between H3K9me3 MtSig and G2/M checkpoint (Fig. 3E), and a negative correlation (R=-0.16, P < 0.001) with immune responses (Fig. 3F). These observations were validated in TCGA data, showing a robust correlation between H3K9me3 MtSig and both G2/M checkpoint and immune responses across diverse cancers (Fig. 4A, B, and Fig. S2). H3K9me3 MtSig displayed positive enrichment for hallmark gene sets related to G2/M checkpoint and oxidative phosphorylation across multiple cancer types. Conversely, it exhibited negative enrichment for immune response pathways, including inflammatory response, interferon alpha response, and interferon gamma response (Fig. S3A). Detailed analyses confirmed these results (Fig. S3B - E).

**Fig. 3.**
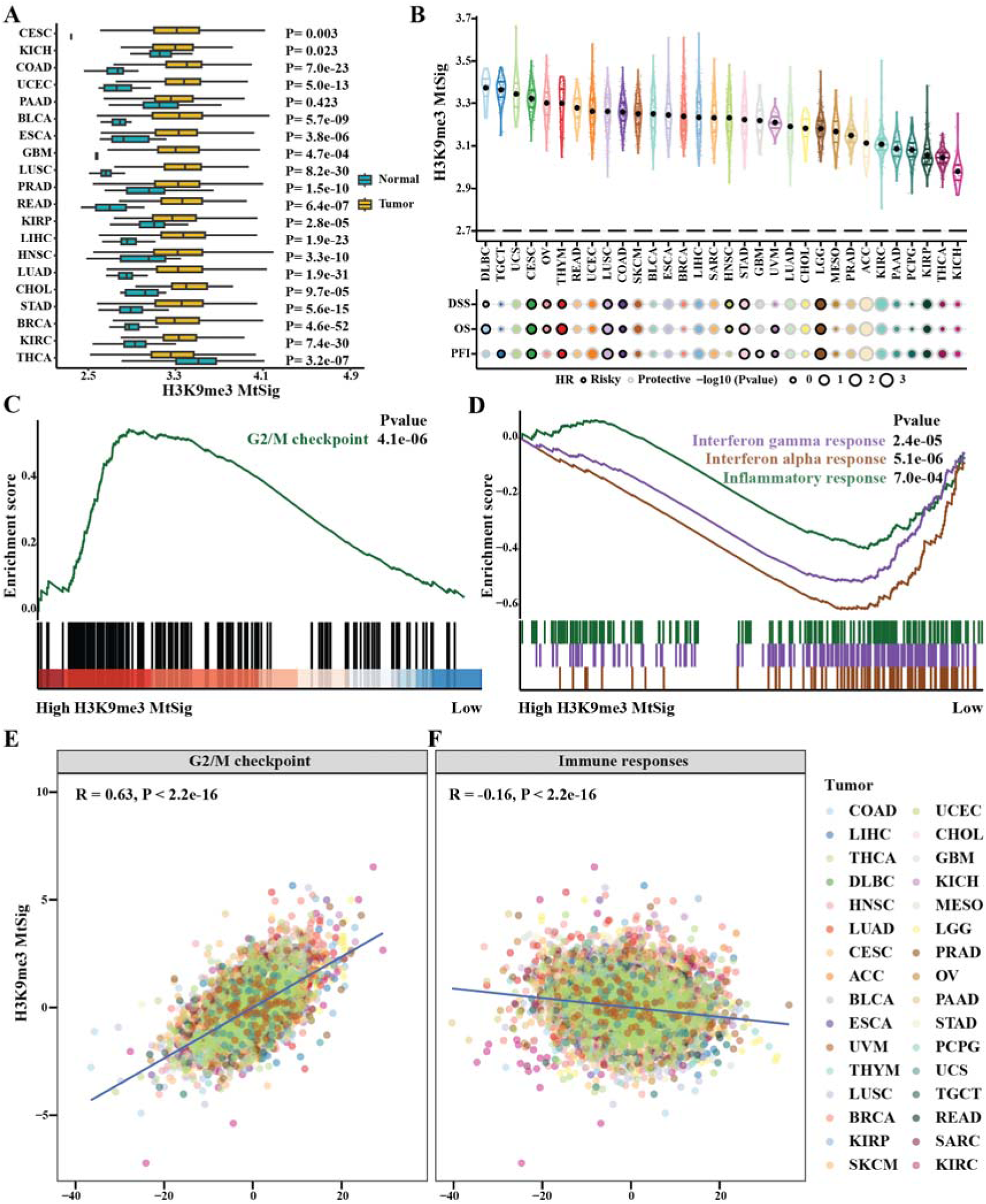
H3K9me3 MtSig and regulated pathways in pan-cancer. (A) The differences in H3K9me3 MtSig between tumor and normal tissues are illustrated. (B) Overall cancer prognosis based on H3K9me3 MtSig score. (C and D) GSEA on hallmark pathways in high versus low H3K9me3 MtSig tumors in pan-cancer. Panel (C) illustrates pathway associated with the G2/M checkpoint, while panel (D) showcases pathways related to immune responses. (E and F) H3K9me3 MtSig is associated with G2/M checkpoint (E) and immune responses (F) in various cancer types.

**Fig. 4.**
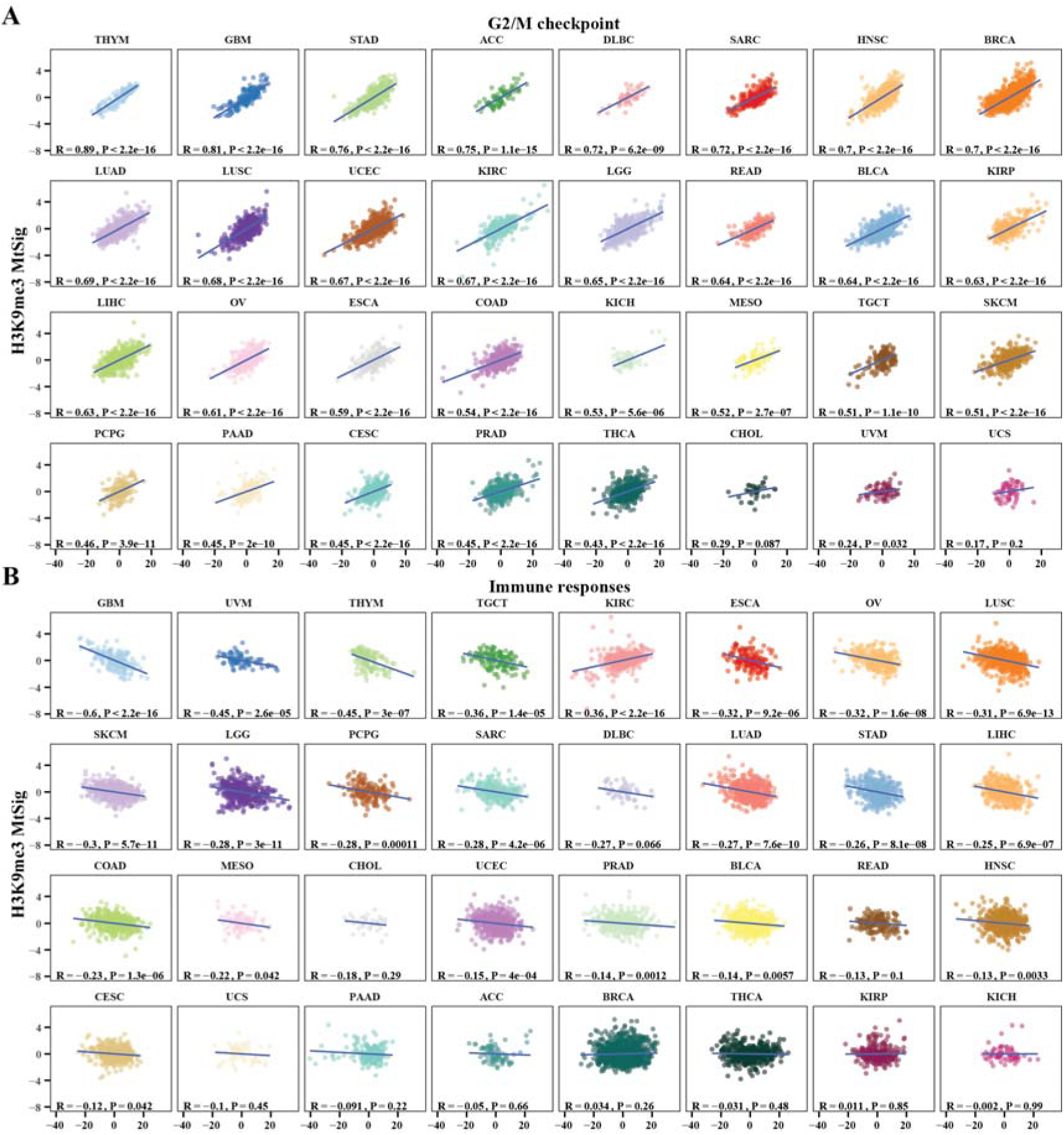
Correlation analysis of H3K9me3 MtSig and two signaling pathways in various cancers. (A and B) The relationship between H3K9me3 MtSig and G2/M checkpoint (A) as well as immune responses (B) is displayed across different types of cancers. The correlation values are notably high in GBM.

### H3K9me3 MtSig regulates repetitive sequence silencing in GBM

The most significant upregulation and prognostic relevance of H3K9me3 MtSig were observed in GBM, LIHC, and LUAD (Fig. 1A, D and Fig. 3A). Given our lab’s focus on GBM, we specifically investigated the role of H3K9me3 MtSig in GBM tumorigenesis and progression. Western blot results from patient-derived GBM cells showed higher expression of SUV39H1, SUV39H2, and SETDB1 compared to normal brain cells (Fig. 5A). IHC staining of GBM tissues and non-neoplastic tissues (Table 1) showed that SUV39H1, SUV39H2, and SETDB1 were upregulated in the GBM tissues (Fig. 5B). Furthermore, interrogating multiple GBM datasets showed that H3K9me3 MtSig was closely associated with the poor survival prognosis including OS, DSS, and progression-free survival (PFS) (Fig. 5C). We found a strong correlation between H3K9me3 MtSig and G2/M checkpoint (R=0.81, P < 0.001) (Fig. 4A), as well as immune responses (R=-0.6, P < 0.001) in GBM (Fig. 4B). Corroboratively, single-cell analysis of H3K9me3 MtSig expression in patient GBM tissues revealed a pattern related to the G2/M cell cycle and showed a negative correlation with immune responses (Fig. 6A - C). H3K9me3 MtSig members correlated positively with key G2/M checkpoint molecules, CDK1, and MKI67 (Fig. 6D and Fig. S4A), while showing a negative correlation with immune responses-associated molecules in GBM (Fig. 6E and Fig. S4B).

**Fig. 5.**
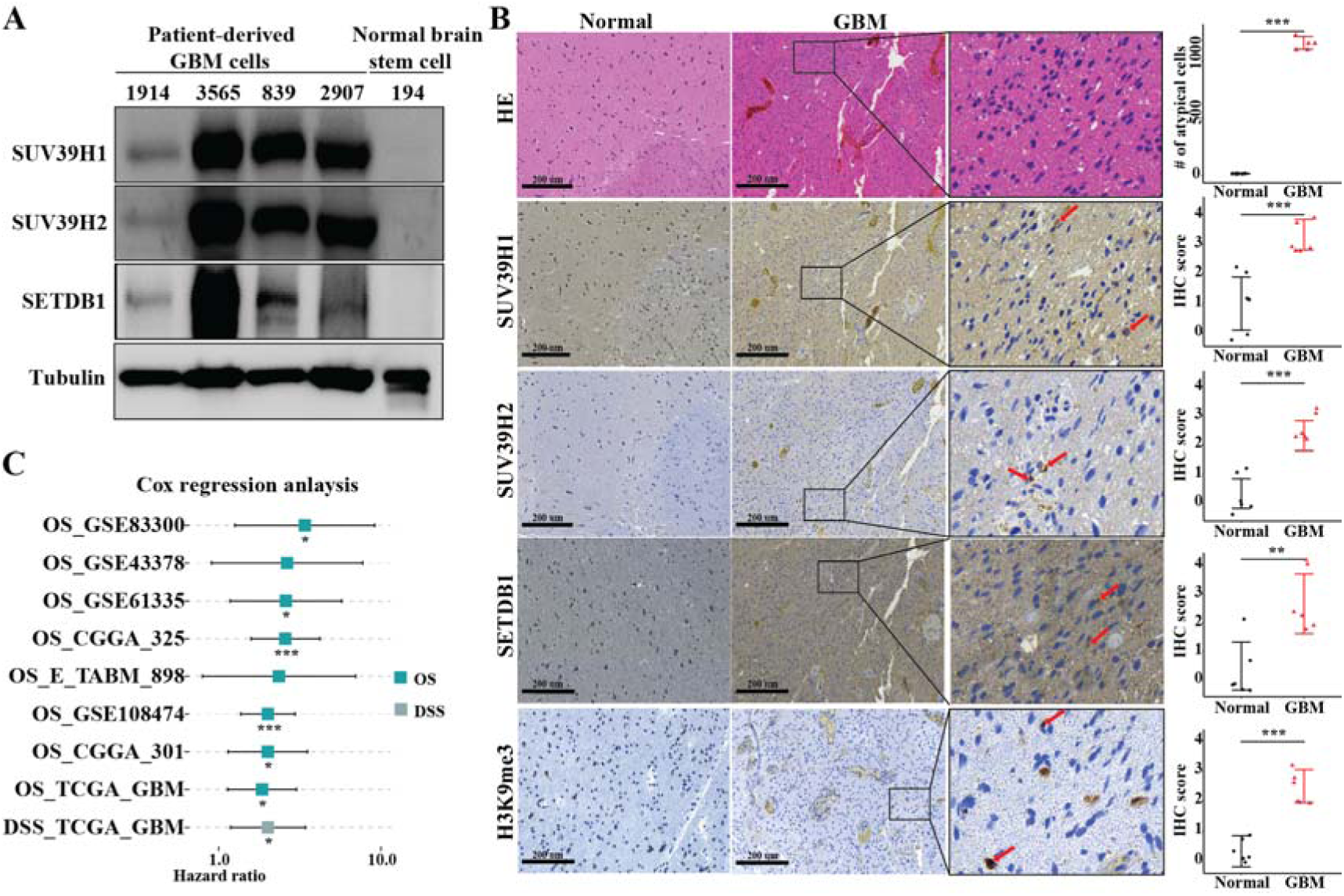
Expression and prognostic relevance of H3K9me3 MtSig in GBM. (A) Western blot showing the expressions of SUV39H1, SUV39H2, SETDB1 in GBM cells and normal brain stem cells. (B) Differential expressions of SUV39H1, SUV39H2, SETDB1 in GBM and non-neoplastic tissues, with representative images (left) and quantification (right). Experiments were performed in triplicate (n=3) and statistical significance was determined using Student’s t-test. **P < 0.01, ***P < 0.001. (C) Forest plot depicting the association between the H3K9me3 MtSig and survival outcomes (OS, DSS) in GBM patients across multiple cohorts. Each row represents a different GBM dataset, with horizontal lines indicating the hazard ratio and 95% confidence interval. A hazard ratio exceeding 1 suggests that higher H3K9me3 MtSig scores correlate with poorer survival outcomes.

**Fig. 6.**
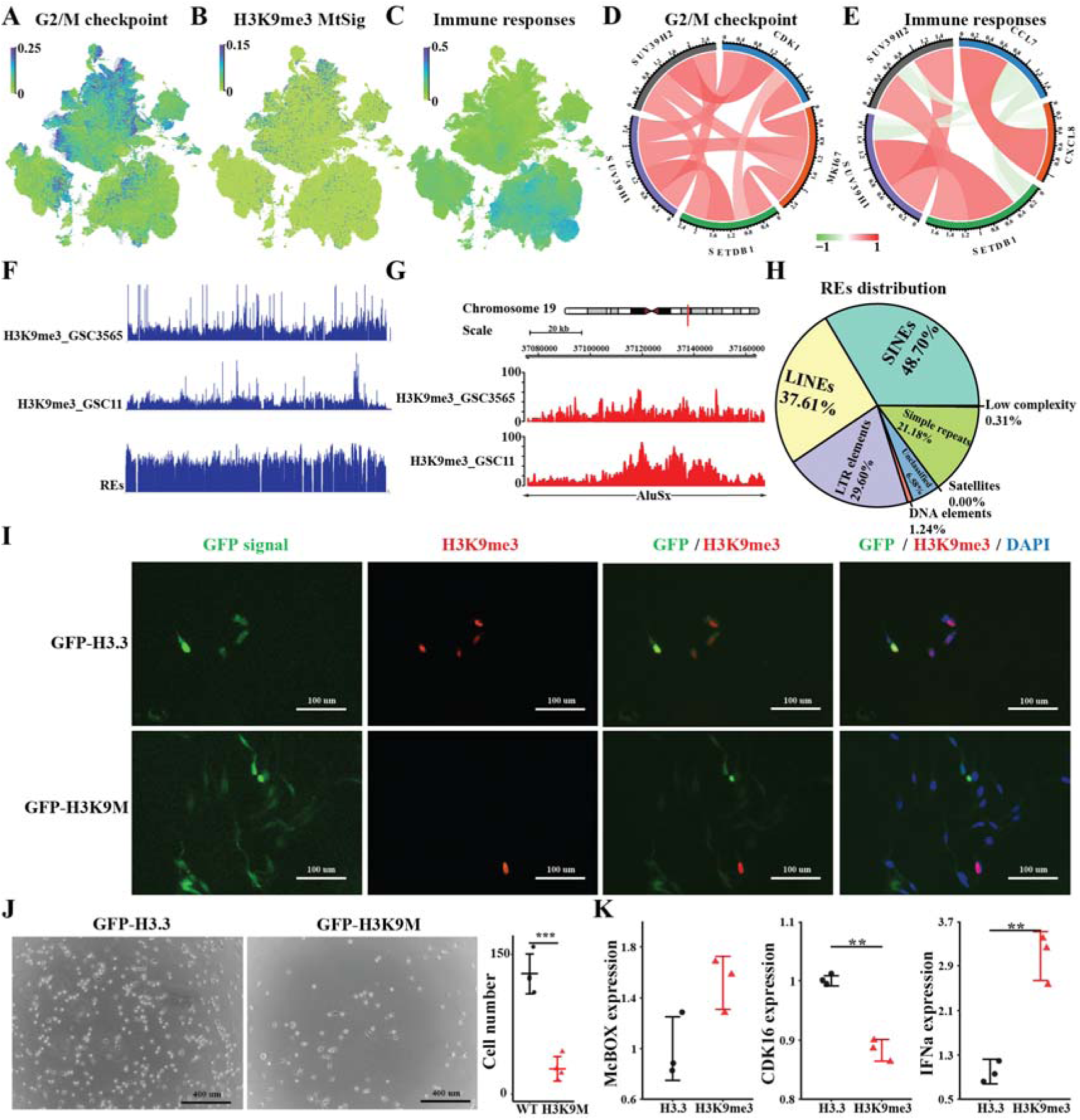
H3K9me3 MtSig regulates G2/M checkpoint and immune responses through H3K9me3-mediated repeat elements (REs) silencing in GBM. (A to C) UMAP visualization of G2/M checkpoint (A), H3K9me3 MtSig (B), and immune responses (C) in cancer cells from GBM tissues by single-cell analysis. (D and E) Analysis of expression correlation between H3K9me3 MtSig genes with *CDK1* and *MKI67* (D), and *CCL7* and *CXCL8* (E) in TCGA GBM datasets. (F and G) Genomic tracks displaying the H3K9me3 ChIP-seq peaks and REs across the entire genome (F), along with a representative RE - AluSx region (G) in GSC3565 and GSC11 cells. (H) RE annotation of H3K9me3 chromatin binding peaks in GSC3565 cells. (I) Immunofluorescence data showing the repression of H3K9me3 by GFP-H3K9M in GSC3565 cells. (J) GFP-H3K9M expression decreases GSC3565 cell proliferation, with representative images (left) and quantification (right). (K) qPCR analysis of McBOX, CDK16, and IFNa expression in GSC3565 cells following indicated transfection. For (J) and (K), data are presented as mean ± SEM (n=3). Statistical analysis was performed using Student’s t-test. **P < 0.01, ***P < 0.001.

We then interrogated how H3K9me3 MtSig regulates these pathways in GBM. Our previous research revealed that H3K9me3, produced by SUV39H1, SUV39H2, and SETDB1 methyltransferases, accumulates in repetitive sequence regions in breast cancer cells (3). We conducted H3K9me3 ChIP-seq in patient-derived GBM cells, demonstrating high enrichment at these repetitive sequences (Fig. 6F, G and Fig. S4C - E), supported by analysis of another H3K9me3 ChIP-seq data from public GBM cell dataset (Fig. 6F, G). These repetitive sequences include short interspersed nuclear elements (SINEs), long interspersed nuclear elements (LINEs), and long terminal repeat (LTR) elements (Fig. 6H). Studies suggest an inverse correlation between repetitive sequence expression and cell cycle progression (33,34) and a positive correlation with immune responses (35,36). We hypothesized H3K9me3 MtSig regulates the G2/M cell cycle and immune responses in GBM via H3K9me3-mediated silencing of repetitive sequences. To test our hypothesis, we depleted the H3K9me3 modification in GBM cells through expressing H3K9M plasmids (37) (Fig. 6I). Depletion of H3K9me3 reduced GBM cell proliferation in comparison to the control (Fig. 6J), accompanied by induction of repetitive sequence mcBox, immune response regulator IFN-α, and downregulation of the G2/M cell cycle regulator CDK16 (Fig. 6K). In summary, these data demonstrate that H3K9me3 MtSig regulates cellular G2/M checkpoint and immune responses through H3K9me-mediated repetitive sequence silencing, contributing to GBM development and poor survival prognosis.

### Clinical prediction model incorporating H3K9me3 MtSig for GBM

To enhance prognostic predictability for adverse events, we devised a comprehensive network score curve utilizing TCGA GBM data. This curve integrates H3K9me3 MtSig with various clinical-pathological factors, including patient age, gender, race, and radiotherapy status (Fig. 7A). Our findings underscore the close relationship between H3K9me3 MtSig and patient prognosis, particularly in the context of radiotherapy, as illustrated by the survival factor association plot (Fig. 7B). Calibration curves for 1-year, 2-year, and 3-year survival rates closely aligned with the standard curve, affirming the accuracy of our network score curve in approximating actual survival probabilities (Fig. 7C). The time-dependent concordance index demonstrated strong model consistency when combining H3K9me3 MtSig with clinical prognostic factors (Fig. 7D). Furthermore, we conducted a revalidation of this clinical predication model using CGGA GBM data, affirming H3K9me3 MtSig as an independent prognostic factor with significant clinical relevance for GBM (Fig. S5).

**Fig. 7.**
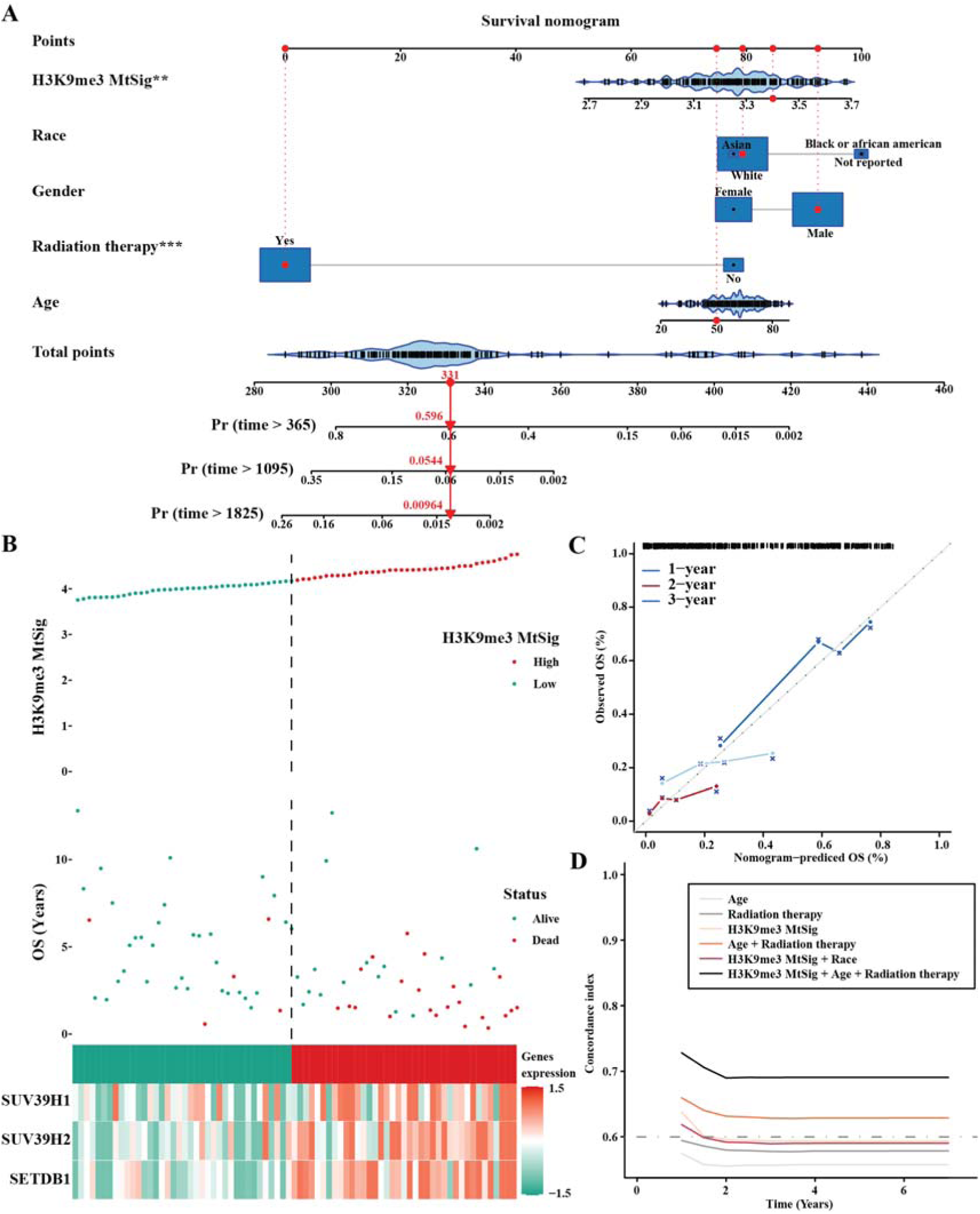
Clinical prediction model incorporating H3K9me3 MtSig using TCGA GBM dataset. (A) Nomogram predicting mortality rate of patients based on clinical-pathological factors and prognostic H3K9me3 MtSig. (B) Risk survival status plot of H3K9me3 MtSig. (C) Calibration of the nomogram for predicting 1-year, 2-year, and 3-year outcomes, showing the consistency between predicted and observed results. The position relative to the 45-degree line indicates the performance of the nomogram, where the 45-degree line represents the ideal prediction. (D) Concordance index showing measure of concordance of predictor with survival of patients.

### Drug susceptibility screening integrating H3K9me3 MtSig for GBM

In our pursuit to assess GBM’s response to diverse drugs or treatments, we employed drug susceptibility screening integrating H3K9me3 MtSig utilizing two methodologies: CTRP2.0 and PRISM. Analysis with CTRP2.0 revealed a notable increase in drug sensitivity, particularly for compounds such as GDC-0879, vemurafenib, MK-0752, BRD-K33199242, and PLX-4720, all showing a significant positive correlation with elevated H3K9me3 MtSig scores (Fig. 8A). Similarly, PRISM analysis demonstrated enhanced drug sensitivity, especially for EPZ-5676, oxymatrine, ganetespib, rubitecan, dirithromycin, and simvastatin, all exhibiting a significant positive correlation with elevated scores (Fig. 8B). These findings underscore the potential of H3K9me3 MtSig as a predictive biomarker and therapeutics for GBM.

**Fig. 8.**
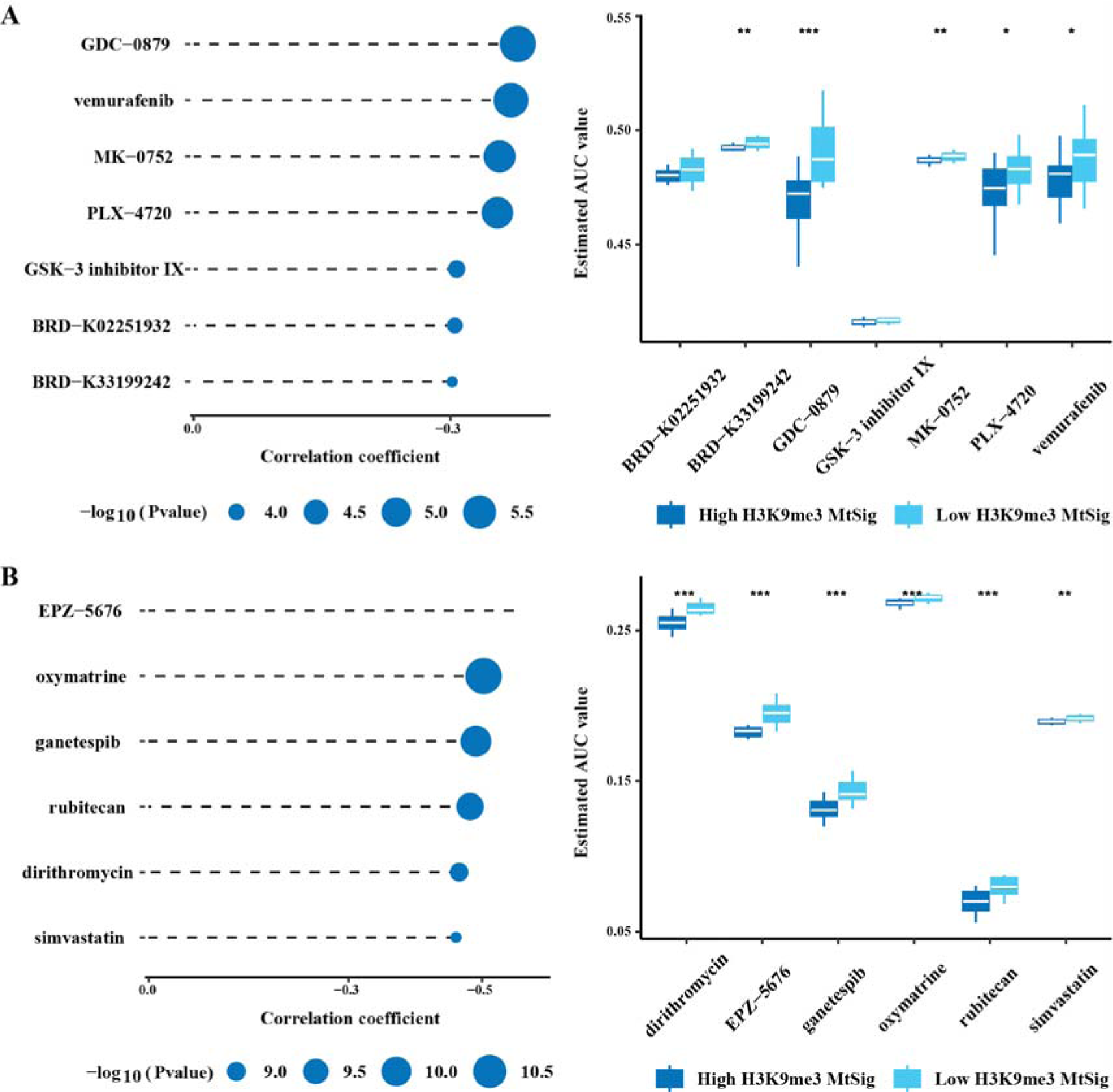
Drug susceptibility screening integrating H3K9me3 MtSig for GBM. (A and B) Analysis of the correlation between H3K9me3 MtSig and drug sensitivity using CTRP2.0 (A) and PRISM (B). GDC-0879, vemurafenib, MK-0752, BRD-K33199242, PLX-4720, EPZ-5676, oxymatrine, ganetespib, rubitecan, dirithromycin, and simvastatin exhibit significant positive correlations with elevated H3K9me3 MtSig.

## Discussion

Our analysis revealed robust correlations of expressions among specific H3K9 methyltransferases across cancers, implying their cooperative interactions. Indeed, studies showed SUV39H1 acts with SETDB1 to suppress developmental genes like *Hox*, accelerating melanoma progression (10). EHMT1/2, SETDB1, and SUV39H1 form a shared megacomplex, maintaining stem cell pluripotency and targeting genes. Depletion of SUV39H1 or EHMT2 destabilizes other methyltransferases at the protein level, highlighting their mutual interdependence (38). These findings suggest mutual regulation among H3K9 methyltransferases, impacting tumorigenesis and progression. We then investigated genetic and epigenetic changes in H3K9 methyltransferase genes. The observed high mutation rates and CNVs in *SUV39H1* and *SUV39H2* may contribute to their functional impairment, while abnormal DNA methylation patterns could influence the expression and activity of *SETDB1* and *SUV39H2*, further compounding their effects on chromatin regulation and oncogenic processes. Considering the intricate interplay among these methyltransferases and their cooperative complex formation, dysregulation may synergistically affect chromatin organization and gene expression. Future studies elucidating how genetic and epigenetic changes disrupt H3K9 methyltransferase activities and complexes are essential for understanding their roles in various cancers.

The integration of H3K9me3 MtSig as a unified signature in our study provided invaluable insights into its role as a prognostic factor and a regulator of signaling pathways pivotal in cancer biology. Our findings clearly demonstrated the marked upregulation of SUV39H1, SUV39H2, and SETDB1 in patient-derived GBM cells and tissue samples, while high expression of the H3K9me3 MtSig is closely associated with poor survival outcomes in GBM patients (Fig. 9). This observation aligns with recent studies reporting frequent overexpression of these H3K9 methyltransferases in GBM, which correlates with adverse patient prognosis (39,40). Beyond its association with the G2/M checkpoint, our analysis revealed robust correlations between H3K9me3 MtSig and various immune response pathways, including the inflammatory response, interferon alpha response, and interferon gamma response. Furthermore, previous studies have highlighted the critical involvement of these immune response pathways in orchestrating antitumor immunity. For instance, the inflammatory response pathway influences immune cell infiltration and function within the tumor microenvironment, ultimately impacting tumor progression and treatment outcomes (41,42). Similarly, interferon signaling pathways play diverse roles in promoting antitumor immunity by enhancing the expression of major histocompatibility complex (MHC) molecules, stimulating immune cell activation, and inducing apoptosis in tumor cells (43–45).

**Fig. 9.**
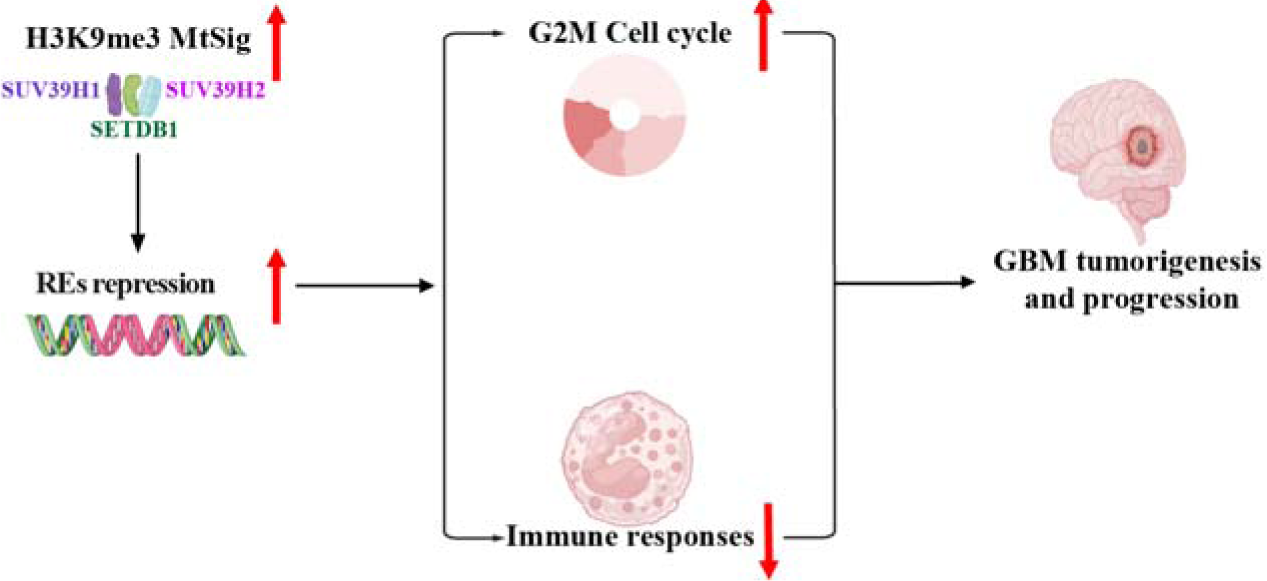
The working model showing the signal cascade between H3K9me3 MtSig, H3K9me3, REs, cell cycle regulation (G2M checkpoint), and immune responses in the context of GBM.

Targeting specific H3K9 methyltransferases, such as SUV39H1 and SETDB1, has been shown to enhance immunotherapy responses in various cancer models. For example, SUV39H1 targeting activates interferon signaling pathways in breast cancer cells, leading to tumor growth arrest and improved response to anti-PD-1 immunotherapy (3). SETDB1 targeting can promote cytotoxic T cell-based immunotherapy in cancer mouse models (46). In the current study, we further validate the critical role of H3K9me3-mediated silencing of repetitive sequences by H3K9me3 methyltransferases in driving these processes by using ChIP-seq experiments. We show significant enrichment of H3K9me3 in repetitive sequence regions, such as SINEs, LINEs, and LTR elements, in GBM cell lines, consistent with our analysis of other public H3K9me3 ChIP-seq data. Importantly, depletion of H3K9me3 in GBM cells resulted in the upregulation of repetitive sequence expression, downregulation of the cell cycle, and induction of the immune responses. Intriguingly, a prior study uncovered the presence of tumor-specific antigens originating from certain repetitive sequences in various cancers, potentially eliciting immune responses (47). These findings offer compelling experimental evidence supporting our hypothesis that the H3K9me3 MtSig regulates cancer cell cycle and immune responses through H3K9me3-mediated silencing of repetitive sequences. Notably, recent studies highlight an intriguing role for SUV39H1 beyond cancer cells, such as regulating chimeric antigen receptor (CAR) T cell signaling for cancer immunotherapy. SUV39H1 targeting enhances the persistence and antitumor efficacy of CAR T cells in lung and disseminated solid tumor models (48), and leukemia and prostate cancer models (49). The intricate interplay between H3K9 methylation, immune response pathways and the tumor microenvironment warrant further investigations to uncover the detailed mechanisms underlying these interactions in GBM and other cancers as such studies may support the development of novel combinatorial therapeutic strategies harnessing both epigenetic modulation and immunotherapy interventions for cancer.

By integrating the H3K9me3 MtSig with clinical-pathological factors, such as age, gender, race, and radiotherapy status, we developed a comprehensive network score curve model, offering compelling insights for incorporating the H3K9me3 MtSig into risk stratification and personalized treatment decisions. Drug susceptibility screening revealed a significant positive correlation between elevated H3K9me3 MtSig and increased sensitivity to various compounds. Notably, several of these compounds have shown promising anti-tumor activities in GBM or other cancers. For example, vemurafenib, a BRAF inhibitor, has demonstrated efficacy in BRAF V600E-mutant GBM patients and is undergoing clinical investigation (50). PLX-4720, another BRAF inhibitor, exhibits potent anti-proliferative effects in GBM cell lines and xenograft models (51). Simvastatin, a cholesterol-lowering drug, displays anti-tumor effects in GBM by inducing apoptosis and inhibiting cell migration and invasion (52). H3K9me3 MtSig could serve as a predictive biomarker for identifying GBM patients likely to respond favorably to targeted therapies or drug repurposing strategies.

The role of H3K9 methyltransferases in various cancers is obviously complex. By using our novel H3K9me3 MtSig based on SUV39H1, SUV39H2, and SETDB1, we can elucidate their potential as prognostic markers, therapeutic targets, and predictors of treatment response across various cancer types. Further, with particular relevance to GBM, our study offers new avenues for clinical intervention.

## Authors’ Contributions

**Q. Xie**: Conceptualization, investigation, visualization, methodology, writing–original draft. **S. Ghosh:** Formal analysis, investigation, writing–original draft. **Y. Du:** Formal analysis, investigation, writing–original draft. **S. Rajendran:** Investigation. **A.A. Cohen-Gadol:** Resources, writing–review and editing. **J.M. Baizabal:** Resources, writing–review and editing. **K.P. Nephew:** Resources, writing–review and editing. **L. Han:** Methodology, writing–review and editing. **J. Shen:** Conceptualization, methodology, funding acquisition, supervision, project administration, writing–original draft, writing–review and editing.

## Acknowledgments

This study received funding from the Indiana University School of Medicine Start-Up Fund (to J. Shen) and the Schwarz Family and Friends Cancer Research Fund (to J. Shen). We thank the ECRO Biorepository at Indiana University Health Methodist Hospital for supplying the normal brain and GBM tissues. Special appreciation to the Histology Core at the Indiana Center for Musculoskeletal Health for their paraffin embedding and sectioning service. We are grateful to Dr. Jeremy Rich at the UPMC Hillman Cancer Center for providing normal brain stem cells and patient-derived GBM cells, and to Dr. Charles Spruck at the Sanford Burnham Prebys Medical Discovery Institute for their research support. Thanks also to Dr. Peter Hollenhorst, Dr. Richard Carpenter, and Dr. Heather M. O’Hagan at Indiana University for their generous contributions in sharing reagents, equipment, and valuable insights during the study.

## Notes

### Competing Interest Statement

The authors have declared no competing interest.

