## Supplementary material for "Multi-omic Analysis Identifies Glioblastoma Dependency on H3K9me3 Methyltransferase Activity": Fig.S1-5

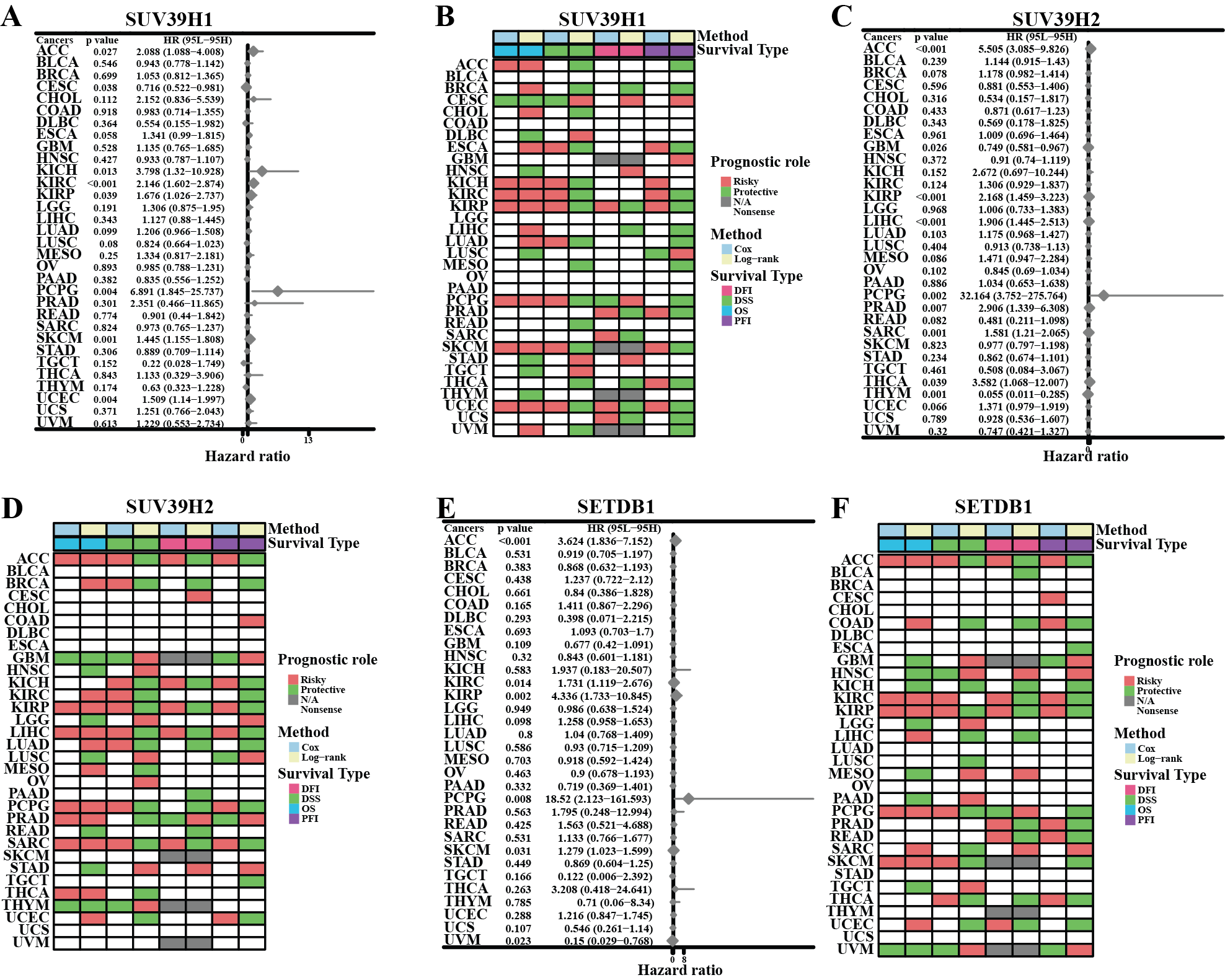

**Fig. S1. Prognostic significance of SUV39H1, SUV39H2, and SETDB1 expression in cancers.** (A, C, E) Summary of the correlations between gene expression and overall survival (OS), disease-specific survival (DSS), disease-free interval (DFI), and progression-free interval (PFI) in cancer patients, based on univariate Cox regression and Kaplan-Meier analysis. Red indicates significant risk factors, and green indicates protective factors (P < 0.05). (B, D, F) Forest plots illustrating the prognostic roles of the respective genes in different cancer types using univariate Cox regression. Red bars indicate statistically significant risk factors.

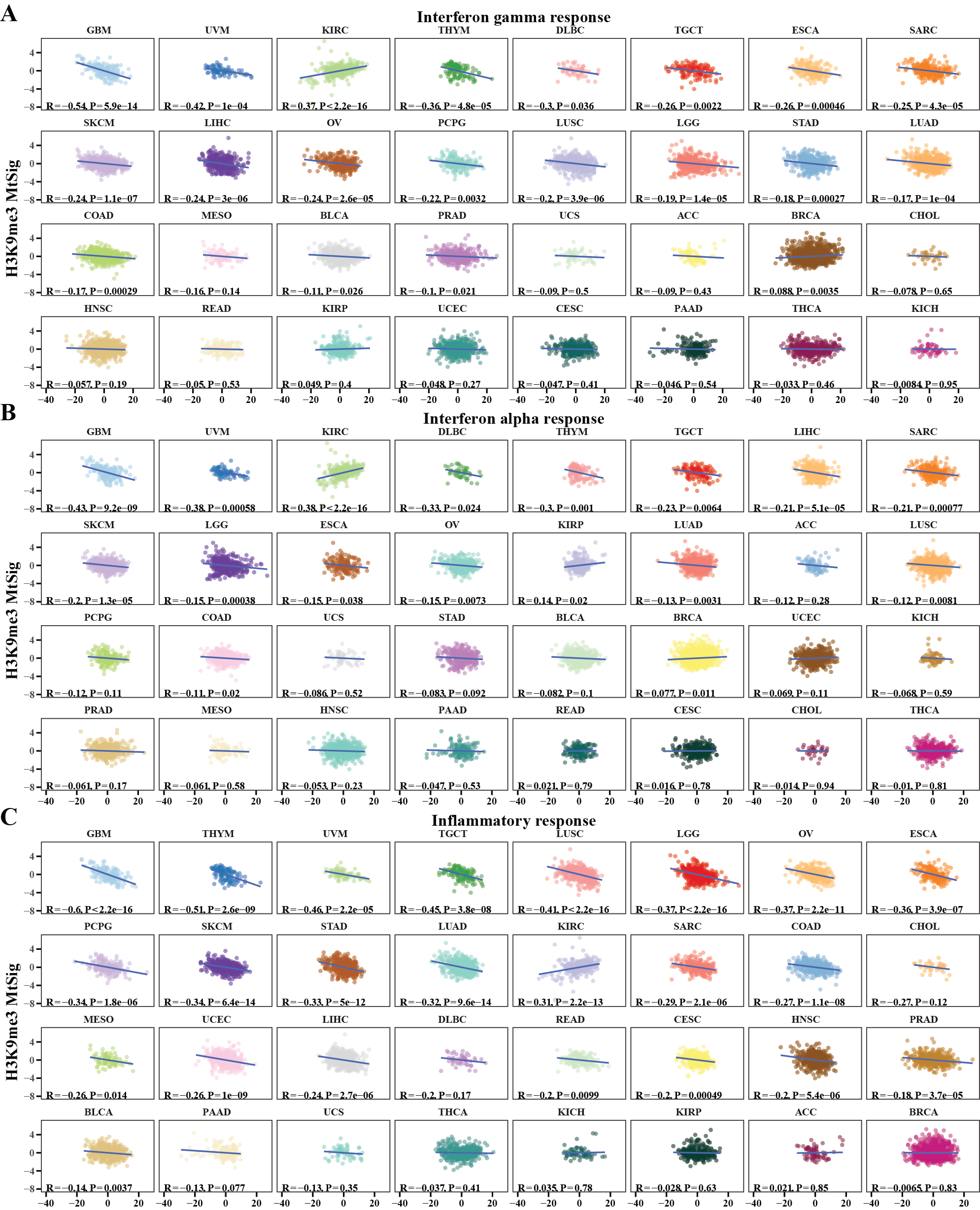

**Fig.S2.** **The correlation between H3K9me3 MtSig and immune responses in pan-cancer.** (A) Analysis of correlations of H3K9me3 MtSig and inflammatory response (A), interferon alpha response (B), and interferon gamma response (C) in various cancers.

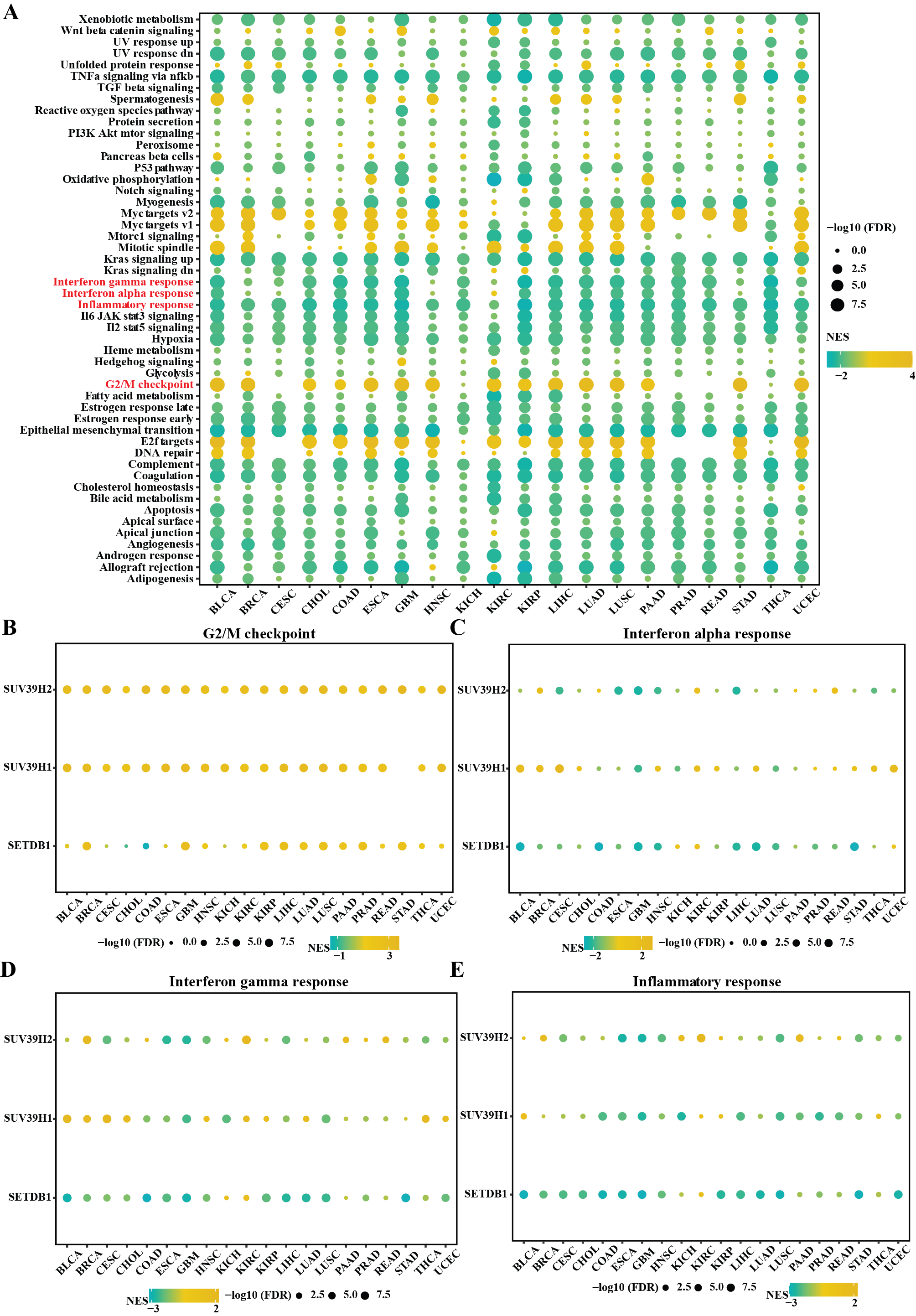

**Fig. S3. Pathways associated with H3K9me3 MtSig across cancers.** (A) Heatmap depicting enrichment scores (normalized enrichment scores, NES) of hallmark gene sets enriched between high and low H3K9me3 MtSig groups across different cancer types by GSEA. (B to E) Analysis of SUV39H1, SUV39H2, and SETDB1 expressions with G2/M checkpoint (B), interferon alpha response (C), interferon gamma response (D), and inflammatory response (E), in various cancer types. Colored bars indicate positive (orange) or negative (blue) correlations.

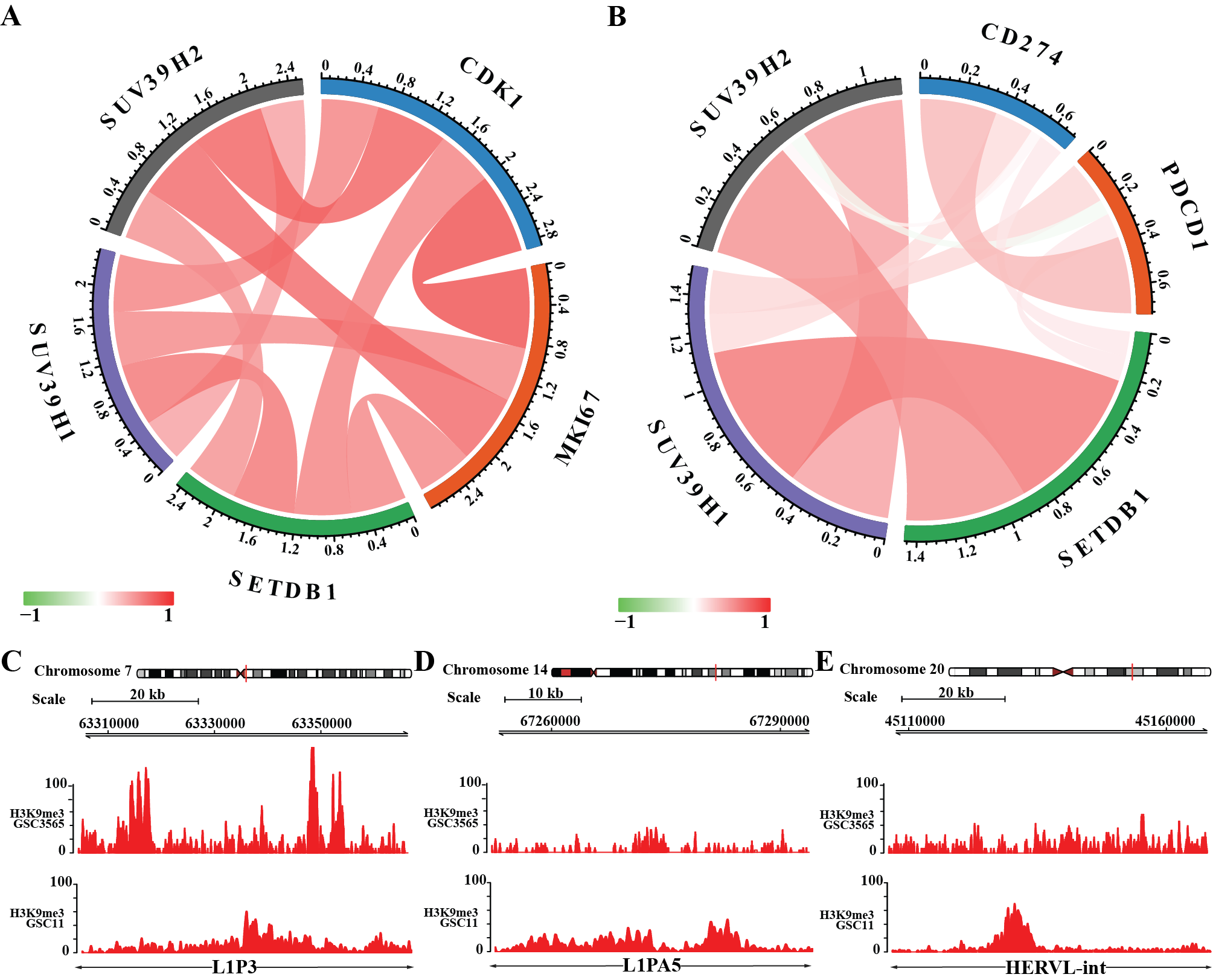

**Fig.S4.** H3K9me3 MtSig regulates G2/M checkpoint and immune responses through H3K9me3-mediated repeat elements (REs) silencing. (A and B) Expression correlations between H3K9me3 MtSig genes with *CDK1* and *MKI67* (A), and *PDCD1* and *CD274* (B) in CGGA GBM database. (C to E) Genomic tracks displaying the H3K9me3 ChIP-seq peaks on three representative REs in GSC3565 and GSC11 cells.

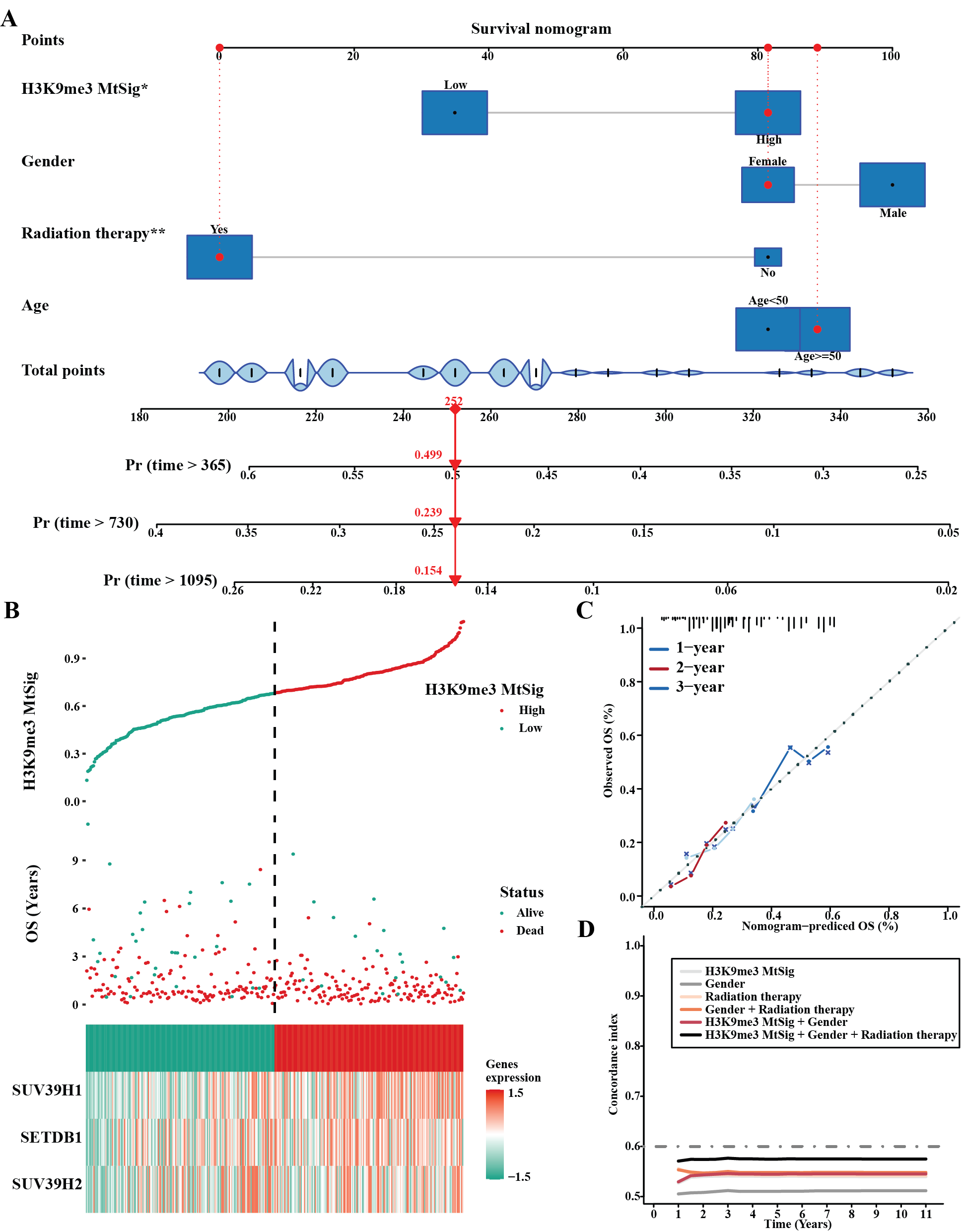

**Fig. S5. Clinical prediction model integrating H3K9me3 MtSig using the CGGA GBM dataset.** (A) Nomogram for predicting patient survival probability by combining H3K9me3 MtSig with clinicopathological factors (gender, age, radiation therapy). The nomogram shows point contributions of each factor towards the total points scale, which maps to predicted survival probabilities. (B) Risk score distribution of H3K9me3 MtSig groups (high vs low) and patient survival status. (C) Calibration curves evaluating the nomogram's performance in predicting 1-year, 2-year, and 3-year survival, by comparing predicted and observed outcomes. Ideal prediction follows the 45-degree line. (D) Concordance index quantifying the predictive accuracy of the model factors (H3K9me3 MtSig, clinicopathological variables) for patient survival times.

**Table S1. TCGA Cancer Types Abbreviations.**

**Table S2. Information for 3 normal and 3 GBM tissues.**
